# FinaleToolkit: Accelerating Cell-Free DNA Fragmentation Analysis with a High-Speed Computational Toolkit

**DOI:** 10.1101/2024.05.29.596414

**Authors:** J.W. Li, R. Bandaru, K. Baliga, Y. Liu

## Abstract

**Motivation:** Cell-free DNA (cfDNA) fragmentation pattern represents a promising non-invasive biomarker for disease diagnosis and prognosis. Numerous fragmentation features, such as end motif and window protection score (WPS), have been characterized in cfDNA genomic sequencing. However, the analytical tools developed in these studies are often not released to the liquid biopsy community or are inefficient for genome-wide analysis in large datasets.

**Results:** To address this gap, we have developed FinaleToolkit, a fast and memory-efficient Python package designed to generate comprehensive fragmentation features from large cfDNA genomic sequencing data. For instance, FinaleToolkit can generate genome-wide WPS features from a ∼100X cfDNA whole-genome sequencing (WGS) dataset with over 1 billion fragments in 0.7 hours, offering up to a ∼50-fold increase in processing speed compared to original implementations in the same dataset. We have bench-marked FinaleToolkit against original approaches or implementations where possible, confirming its efficacy. Furthermore, FinaleToolkit enabled the genome-wide analysis of fragmentation patterns over arbitrary genomic intervals, significantly boosting the performance for cancer early detection.

**Availability and implementation:** FinaleToolkit is open source and thoroughly documented with both command line interface and Python application programming interface (API) to facilitate its wide-spread adoption and use within the research community: https://github.com/epifluidlab/FinaleToolkit.

## 1 Introduction

Circulating cell-free DNA (cfDNA) in peripheral blood has been widely utilized as a noninvasive biomarker for cancer diagnosis and prognosis (Snyder *et al*. 2016; Cristiano *et al*. 2019; Jiang *et al*. 2020; Guo *et al*. 2022). cfDNA is characterized by highly nonuniform fragmentation across the genome, with patterns that have been linked to various epigenetic marks within the originating cells (Snyder *et al*. 2016; Zhou *et al*. 2022a, 2022b). Numerous fragmentation pattern features have been identified in cfDNA genomic sequencing, including both raw and adjusted window protection scores (WPS) (Snyder *et al*. 2016), DNA evaluation of fragments for early interception (DELFI) (Cristiano *et al*. 2019), end motifs, motif diversity scores (MDS) (Jiang *et al*. 2020), breakpoint motifs (Guo *et al*. 2022), and cleavage ratios near CpG sites (Zhou *et al*. 2022a). These patterns have been shown as promising biomarkers to boost the performance of cfDNA for disease diagnosis and prognosis (Cristiano *et al*. 2019). However, source codes from these studies are often absent, poorly documented, non-executable, or not ready for the genome-wide analysis of large genomic datasets, which poses a significant barrier to the clinical application of cfDNA fragmentomics.

Here, we introduce FinaleToolkit (**Fig.1**), a standalone, open-source, and thoroughly documented Python package that can efficiently extract over ten popular fragmentation patterns from a large cfDNA genomic sequencing dataset. FinaleToolkit replicates fragmentation patterns previously published, facilitating analyses when original source codes are unavailable or impractical to use. This tool supports parallel processing for all the features, allowing for the processing of large genomic datasets with a speed increase of up to 50 times compared to existing implementations. Its rapid and memory-efficient design enables comprehensive genome-wide analysis of various fragmentation features across specified genomic intervals, which substantially boosts the performance of cfDNA fragmentomics for cancer early detection. Existing cfDNA fragmentation analysis end-to-end workflows that accept FASTQs are not easily adapted by users who already possess aligned data or who need modular access to fragmentomic computations. During review of this work, cfDNAPro (Wang *et al*. 2025) was published and can extract features from mapped BAM files; however, it does not deliver comprehensive, high-throughput fragmentation feature extraction at the cohort scale. Therefore, there remains a clear need for a flexible, standalone command-line tool focused specifically on rapid, scalable generation of diverse fragmentomic features.

**Figure 1.**
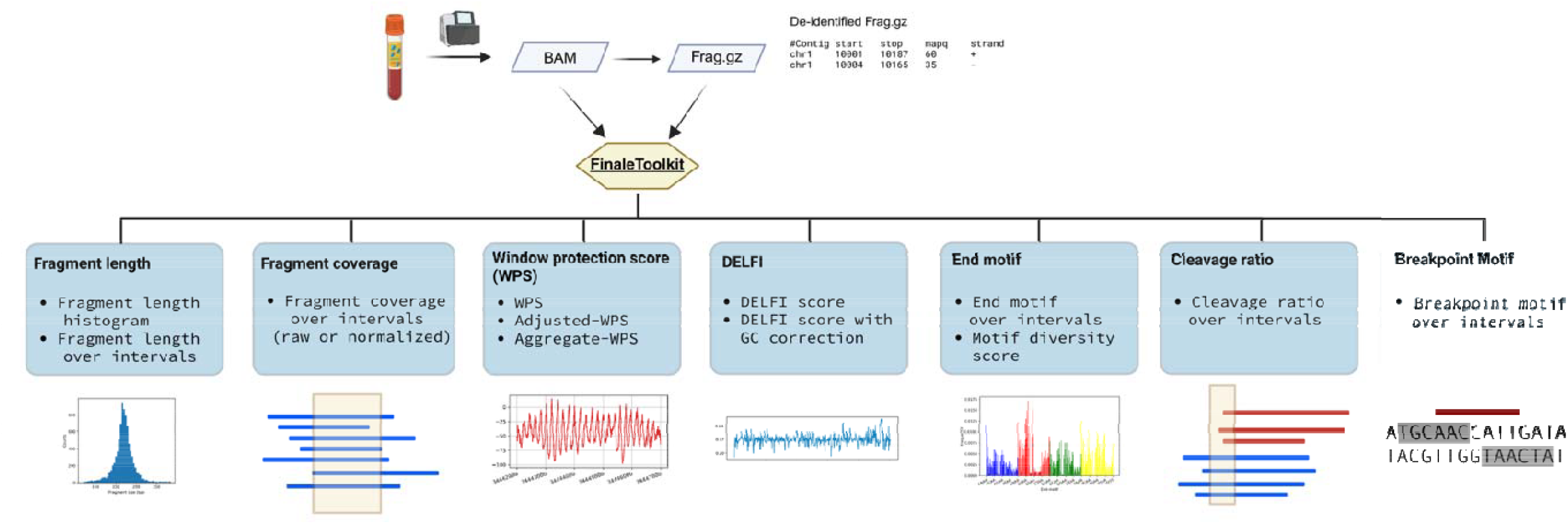
FinaleToolkit workflow. Created with BioRender.com.

## 2 Results

We initially employed publicly available cfDNA deep WGS data (BH01 (Snyder *et al*. 2016), ∼100X, ∼1 billion fragments in total, ∼168Gb bam file) from healthy and non-pregnant individuals as our primary dataset to replicate fragmentation features reported in prior studies. The genome-wide fragment length distribution generated by our tools matches expected cfDNA-specific patterns (**Fig. 2a**), including 167 bp mode and 10.4 bp periodicity (Snyder *et al*. 2016). Similarly, fragment coverage distributions near transcription start sites (TSS) and transcription factor binding sites (TFBS) exhibited the characteristic signatures associated with nucleosome phasing (**Fig. 2b**). Furthermore, genome-wide window protection scores (WPS) demonstrated a high correlation with results from the original implementations (Snyder *et al*. 2016) (**Fig. 2c-f**, Pearson correlation coefficient > 0.99, p-value < 2.2e^-16^).

**Figure 2:**
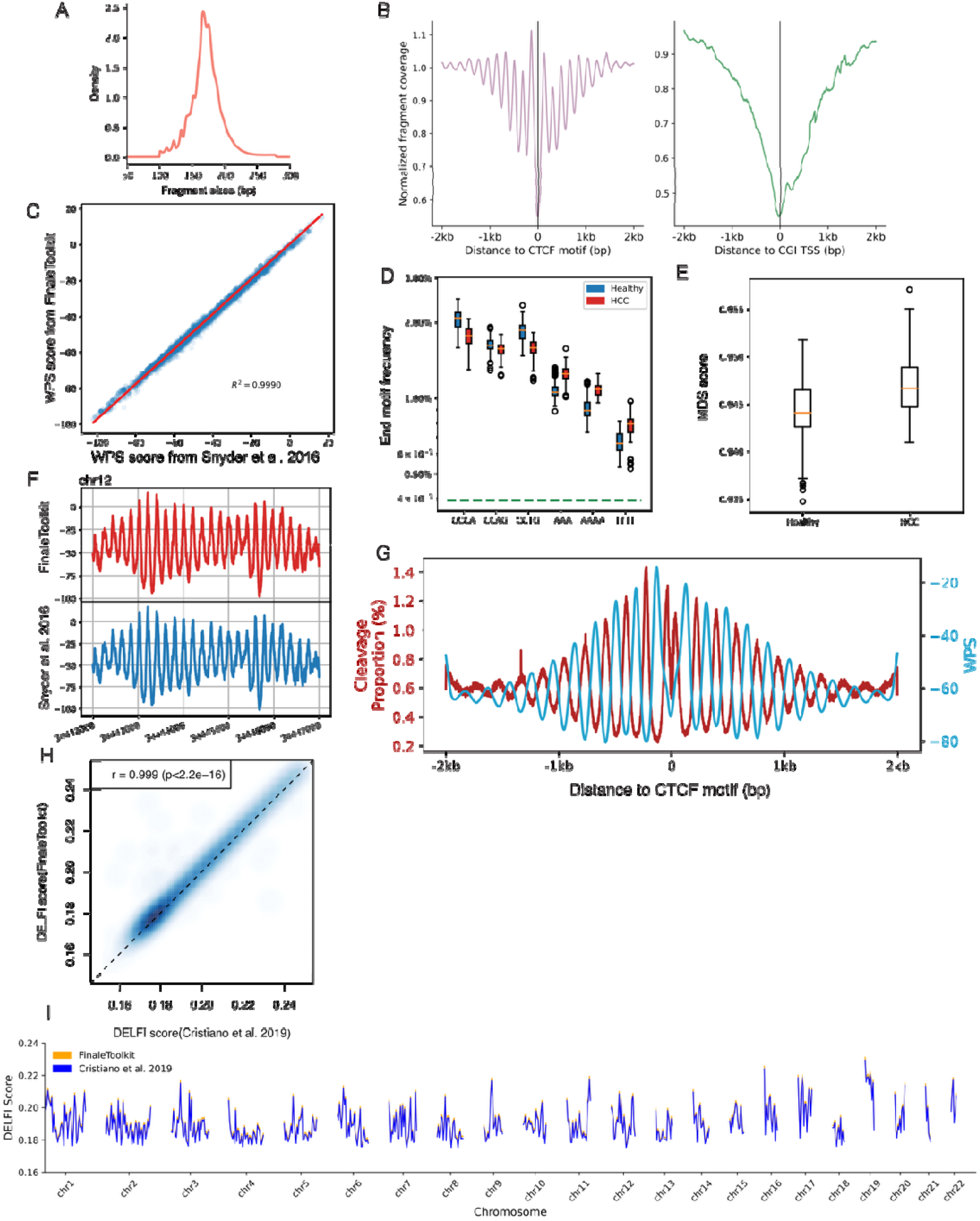
FinaleToolkit generated concordant cfDNA fragmentation patterns similar to those from the previous studies. (a). Histogram of cfDNA fragment size distribution. (b). Normalized fragment coverages near transcriptional starting sites and CTCF motifs. (c) Comparison of genome-wide WPS between FinaleToolkit and Snyder et al.’s 2016 Cell implementation (d). end motif differences between HCC and healthy (e). Boxplot of MDS between HCC and healthy. (f). Example plot of WPS score generated from BH01 using FinaleToolkit (upper panel) and using Snyder et al.’s 2016 Cell (lower panel) at a specific genomic region in chr12. (g). Cleavage ratio and WPS near CTCF motifs. (h) A scatterplot of genome-wide DELFI from BH01 was calculated by FinaleToolkit vs. that calculated from a modified version of DELFI. Pearson correlation coefficient and p-value were calculated using cor.test in R (v4.2.2). (i).Genome-wide DELFI score (chr1-22) generated by modified script from Cristiano et al. 2019 (blue) and FinaleToolkit (orange) at BH01 data.

Due to the original DELFI source code being non-executable (Cristiano *et al*. 2019), we compared FinaleToolkit with a fork of the original DELFI scripts (see Methods in detail) and achieved comparable patterns (**Fig. 2h-i**, Pearson correlation coefficient > 0.99, p-value < 2.2e^-16^). To replicate the differences in end motif and MDS observed between Hepatocellular carcinoma (HCC) patients and healthy individuals, we utilized publicly available cfDNA WGS data from both groups (Jiang *et al*. 2015; Cristiano *et al*. 2019; Zhou *et al*. 2022b). The source code for analyzing end motifs was not provided with the original publication, but an open source implementation called Freefly has been made available by one of the authors: https://github.com/hellosunking/Freefly. When comparing Freefly to Finaletoolkit for calculating end motif frequencies, we found that the results of the two softwares are highly similar (χ^2^=0.00112265). When comparing to the original author’s figures, FinaleToolkit produced similar results for the end motif and MDS despite using a different dataset (**Fig. 2d-e**). A recent study suggested a tight correlation between the cleavage ratios near CpG sites and the DNA methylation status at CpGs (Zhou *et al*. 2022a). Our FinaleToolkit enabled the efficient extraction of the genome-wide cleavage ratios feature from deep cfDNA WGS and observed the anti-correlated phasing patterns between nucleosomes (WPS) and cleavage ratio near CTCF, similar to the previous findings of anti-correlated phasing patterns between nucleosomes and DNA methylation patterns near the same motifs (Kelly *et al*. 2012) (**Fig. 2g**). Overall, FinaleToolkit successfully computed fragmentation patterns consistent with those identified in previous research.

We subsequently benchmarked the processing speed and memory usage of FinaleToolkit against the original source code and other popular implementations when available. For the WPS study, FinaleToolkit processed genome-wide WPS features in 16.6 million windows from BH01 data in 0.70 hours using 64 CPU cores and 40 GB of memory, achieving approximately a 50-fold increase in processing speed compared to the original implementation (**Fig. 3a**). Remarkably, even with a single CPU core, FinaleToolkit delivered results more than twice as fast as the baseline. Although an author’s implementation of DELFI is available, we were unable to adapt this code to use on fragmentation data aligned to GRCh38. By using FinaleToolkit, DELFI could be calculated within 12 minutes (**Fig. 3e**) and 3.3 GB of memory with 16 processes (**Supplementary Figure 1e**). We also assessed the speed and memory efficiency for processing other fragmentation features on the same dataset (**Fig. 3b-d,f**). All remaining features were calculated within 1 hour with 64 CPUs and reasonable memory cost (**Supplementary Figure 1**). It should be noted that FinaleToolkit’s memory efficiency allows it to handle larger data than similar tools. For instance, when we attempted to benchmark the end motif implementation of cfDNApro on BH01, the software exceeded the 256 GB memory limit of our HPC cluster, while Finaletoolkit can calculate genome-wide end motifs with less than 360 MB used (**Supplementary Figure 1d**). Since many of the fragmentation studies were done at low coverage, we also demonstrated the performance at a typical ∼1X cfDNA WGS dataset with ∼20 million high-quality cfDNA fragments. Any given fragmentation feature may be calculated within 40 minutes with 15 GB of memory and 16 CPUs (**Supplementary Figure 2**).

**Figure 3.**
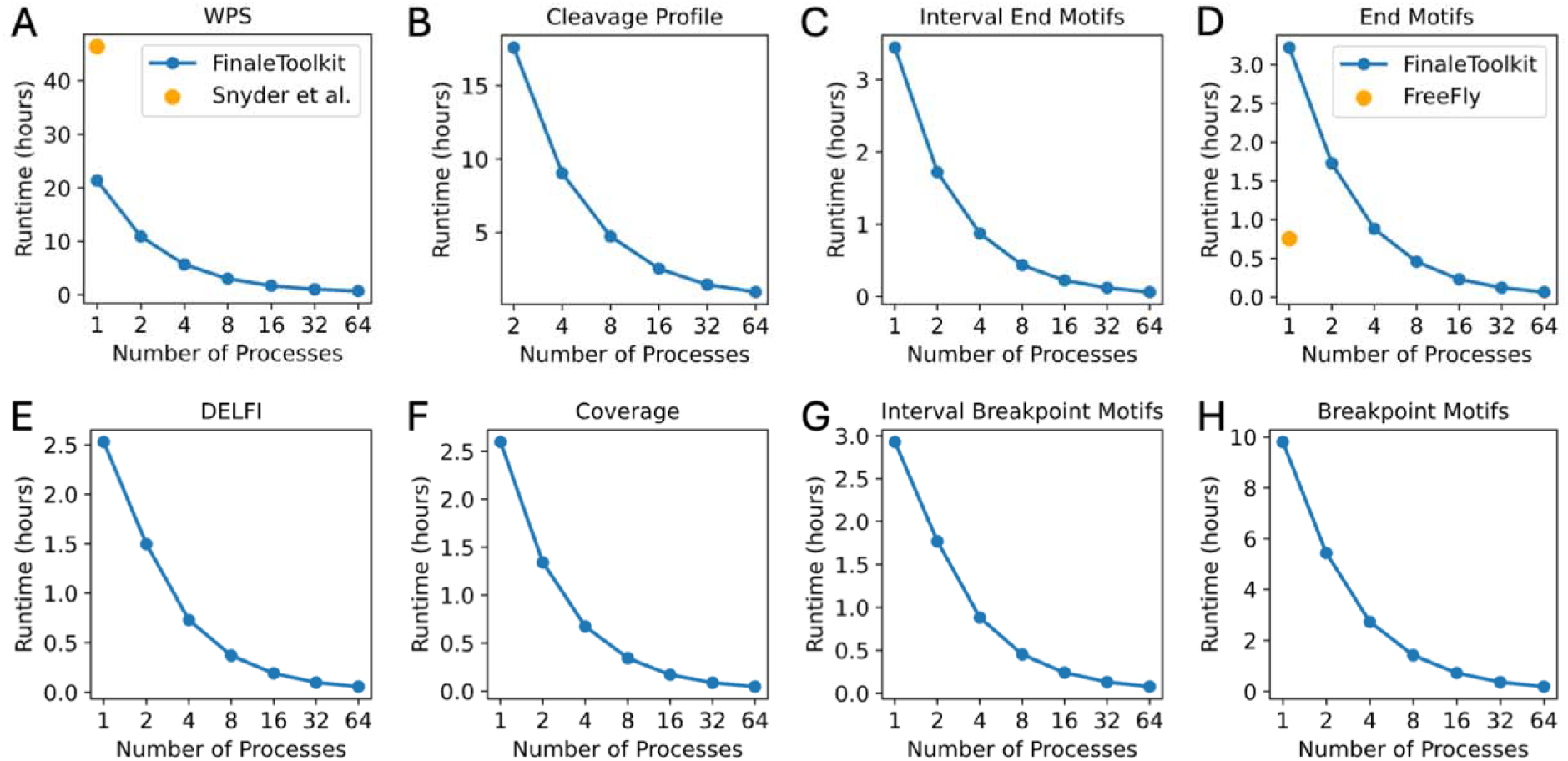
FinaleToolkit demonstrates improved speed for genome-wide analysis of fragmentation patterns from large cfDNA genomic datasets. Wall time cost to calculate (a) WPS vs original implementation, (b) cleavage ratio, (c) end motifs over intervals, (d) genome-wide end motifs vs Freefly implementation, (e) DELFI, (f) fragment coverage, (g) breakpoint motifs over intervals, and (h) breakpoint motifs.

Due to its memory and runtime efficiency, FinaleToolkit has facilitated the analysis of cfDNA fragmentation patterns across arbitrary genomic intervals, including previously unexplored measures such as end motif and MDS over genome-wide non-overlapping intervals. Consequently, we have further demonstrated its utility in identifying aberrations in fragmentation patterns at specific genomic intervals, thereby increasing the predictive power of models trained on these features. A prior study identified significant differences in genome-wide summary statistics for 256 different end motifs and associated single MDS scores between early-stage cancer patients and healthy controls (Jiang *et al*. 2020). We applied this strategy to our previously generated cfDNA WGS dataset from 25 matched pairs of early-stage breast cancer patients and healthy controls (matched 1:1 by age, gender, race, alcohol usage, and smoking history) (Zhou *et al*. 2022b). Using the original approach, classification power was limited, with an Area Under the ROC Curve (AUC) of 0.44 in cross-validation (**Supplementary Figure 3a**). By locating the top 1 or 100 most variable genomic intervals in the training fold, we substantially improved the classification performance on the same data split, achieving an AUC of 0.70 or 0.84, respectively (**Supplementary Figure 3b-c**). In summary, the advanced computational capabilities of FinaleToolkit substantially accelerate genome-wide analysis of fragmentation patterns in large genomic datasets.

## 3 Conclusions and Discussion

In summary, we developed FinaleToolkit to expedite the fragmentomic analysis of large-scale cfDNA genomic sequencing datasets. Current clinical applications of cfDNA fragmentomics frequently necessitate that researchers either reimplement well-known algorithms or modify existing codes for specific requirements. We have verified the efficacy of FinaleToolkit by replicating fragmentation patterns observed in prior studies or benchmarking them with the original code when available. Additionally, genome-wide summary statistics, such as end motif and MDS, ignored the variations of fragmentation patterns at different genomic regions, which may provide additional information to distinguish abnormal disease status. Our toolkit here enables genome-wide fragmentomic analysis at arbitrary genomic intervals through parallel processing, which was not available with original software implementations for large datasets. These additions improve classification performance for separating early-stage cancer patients from matched healthy controls. At present, our focus is limited to popular fragmentation patterns for which source code is unavailable or unsuitable for large datasets. However, FinaleToolkit’s modularity allows for additional features to be progressively integrated into FinaleToolkit in the near future.

## 4 Methods

### 4.1 Ethics approval and consent to Participate

Not applicable.

### 4.2 Data preprocessing of cfDNA WGS data

Raw sequencing datasets were processed by our previously published cfDNA workflow (https://github.com/epifluidlab/finaledb_workflow) (Zheng, Zhu and Liu 2021) and downloaded from FinaleDB (Zheng, Zhu and Liu 2021). The workflow is managed by snakemake v5.19. Specifically in the workflow, raw fastq files were trimmed by Trimmomatic v0.39 and then mapped to the human genome (GRCh37 and GRCh38) using BWA-MEM 0.7.15 with default parameters. Only high-quality reads were considered in the analysis (high quality: uniquely mapped, no PCR duplicates, both ends are mapped with mapping qualities more than 30, and properly paired).

### 4.3 Interface of FinaleToolkit

FinaleToolkit is both a command-line program and a Python library that is accessible from the PyPI package repository using pip, Python’s native package manager, and from Bioconda using the widely used conda package manager. This allows for easy, automated installation. Input consists of sequenced aligned reads (BAM, CRAM) or a FinaleDB style tab-separated, tabix-indexed frag.gz or frag.tsv.bgz. When used in the command line, files can be pipelined into and out of select commands. As FinaleToolkit processes fragment coordinates after pre-processing, its tools exhibit broad compatibility across various sequencing platforms and aligners. A Snakemake workflow is available for ease of use in genomics pipelines.

### 4.4 Framework of FinaleToolkit

To make FinaleToolkit more maintainable, expandable, and efficient, we implemented several functions in the *finaletoolkit*.*utils* module that recurs across multiple features, including functions to read one of several possible fragmentation file formats and return the fragment coordinates functions to filter reads by flags and duplication, and functions to parse BED files.

### 4.5 Benchmarking the speed and memory cost in FinaleToolkit

We calculated the wall time and memory cost with different numbers of CPU cores using a single high-performance computing cluster node (One Intel(R) Xeon(R) Gold 6338 CPU (64 cores) @ 2.0GHz with 256GB DDR4 memory). All benchmarks for the high-coverage dataset were performed using BH01 WGS data. Benchmarking wall time and memory cost were measured using the default Linux *time* command and Slurm logs, respectively. Benchmarks were only for the *finaletoolkit* command for each feature, and do not include any preprocessing steps. Benchmark results may differ the first time scripts are run due to the caching behavior of the HTSlib dependency of pysam. Specific benchmarking details are provided in the supplementary materials.

### 4.6 Classification of early-stage cancers

We utilized our previously published cfDNA WGS dataset from 25 pairs of early-stage breast cancer and matched healthy controls (Zhou *et al*. 2022b). We started with the de-identified fragment files from zenodo.org (breast.tar). To obtain genome-wide MDS, we first ran *finaletoolkit end motifs* and then ran *finaletoolkit mds* on the resulting tab-separated file to generate MDS data for each sample. To obtain MDS at different genomic intervals, we ran *finaletoolkit interval-end motifs* with default parameters on 100kb non-overlapped intervals from autosomes in GRCh37 and then ran *finaletoolkit interval-mds* on the resulting tab-separated file to generate MDS data for each interval in each sample. The samples were split into a ten-fold cross-validation and repeated ten times by using RepeatedStratifiedKFold from scikit-learn(v1.4.2). To validation and repeated ten identify the most variable features from the interval MDS dataset, we employed an ANOVA F-value-based feature selection method. The implementation utilizes the *SelectKBest* module from scikitlearn(v1.4.2), and selects the top *k* features with the highest F-value in the training set. In the end, we utilized *LogisticRegression* from *scikit-learn*(v1.4.2) with the following parameters: “penalty=‘l2’, dual=False, tol=0.0001, C=1.0, fit_intercept=True, intercept_scaling=1, class_weight=None, solver=‘lbfgs’, max_iter=100, multi_class=‘auto’, verbose=0, warm_start=False, l1_ratio=None” to calculate ROC. To make a fair comparison, we generated the same sample split for the study of genome-wide MDS and interval MDS.

## Acknowledgements

This research was supported in part through the computational resources and staff contributions provided by the Genomics Compute Cluster, which is jointly supported by the Feinberg School of Medicine, the Center for Genetic Medicine, and Feinberg’s Department of Biochemistry and Molecular Genetics, the Office of the Provost, the Office for Research, and Northwestern Information Technology. The Genomics Compute Cluster is part of Quest, Northwestern University’s high-performance computing facility, with the purpose to advance research in genomics. This work used Bridges-2 RM and Ocean storage at Pittsburgh Supercomputing Center (PSC) through allocation MCB190124P from the Advanced Cyberinfrastructure Coordination Ecosystem: Services & Support (ACCESS), which is supported by National Science Foundation grants #2138259, #2138286, #2138307, #2137603, and #2138296.

## Supplementary data

Supplementary data are available at *Bioinformatics Advances* online.

## Data availability

The publicly available cfDNA WGS data used in this study are available in the Gene Expression Omnibus (GEO) database under accession code [BH01: GSE71378] and the European Genome–Phenome Archive (EGA) database under accession code [HCC and healthy: EGAS00001001024]. The de-identified fragment files are also available from FinaleDB and Zenodo [breast cancer and healthy (breast.tar): https://zenodo.org/records/6914806].

## Software availability

The code for FinaleToolkit and associated scripts is publicly available on GitHub under the MIT license: https://github.com/epifluidlab/FinaleToolkit. The documentation for FinaleToolkit and related Application Programming Interface (API) is available at: https://epifluidlab.github.io/FinaleToolkit/. Further usage instructions, tutorials, and instructions for replicating figures can be found at: https://github.com/epifluidlab/FinaleToolkit/wiki.

## Contributions

Y.L. conceived the project. J.W.L., R.B., K.B., and Y.L. wrote the code for FinaleToolkit. J.W.L. and R.B. benchmarked the performance. Y.L., J.W.L., and R.B. wrote the manuscript together. All authors read and approved the final manuscript.

