## Supplementary Materials for "FinaleToolkit: Accelerating Cell-Free DNA Fragmentation Analysis with a High-Speed Computational Toolkit"

James Wenhan Li[1-3,*], Ravi Bandaru[1-2,*], Kundan Baliga[4], Yaping Liu[1-2,#]

**Affiliations:**
1. Department of Biochemistry and Molecular Genetics, Feinberg School of Medicine, Northwestern University, Chicago, IL 60611, USA
2. Robert H. Lurie Comprehensive Cancer Center of Northwestern University, Chicago, IL 60611, USA
3. Department of Computer Science, Wake Forest University, Winston-Salem, NC 27109, USA
4. Neuqua Valley High School, Naperville, IL 60564, USA

* Contributed equally as co-first authors
### Corresponding

#### Supplementary Method

##### Benchmark WPS score

The original WPS implementation (Snyder *et al.* 2016) used the pipeline of three programs: an early fork of samtools to read a BAM file, a Python 2 program *FilterUniqueBam.py* to filter reads, and a Python 2 program *extractReadStartsFromBAM2Wig.py*, producing a wig.gz file containing raw WPS scores over a single interval. For benchmarking, we ran this pipeline for every 10 kb interval spanning the whole genome on a ~100x coverage file (BH01). We were unable to run this genome-wide without intervals due to the memory usage exceeding the 256 GB limit of our cluster. When benchmarking FinaleToolkit, we ran *finaletoolkit wps* on BH01 over the whole genome with 1, 2, 4, 8, 16, 32, and 64 worker processes.

Although our results are very similar to that of Snyder et al. ($R^2$>0.999), there still exists a small discrepancy between the results generated by these two programs. This is due to slight differences in how each program filters reads. Snyder et al. use a unique fork of samtools that filters fragments by size, whereas we wrote our own code to perform this task.

##### Benchmark DELFI score

To benchmark FinaleToolkit, we ran *finaletoolkit delfi* on BH01 over 100kb genome-wide bins and then merged these bins into 5Mb windows as in the original study. We used the Encode Data Analysis Center blacklisted regions as a filter, UCSC Genome Browser hg19 (GRCh37) gaps track for centromere and telomere coordinates, and used the options for GC-correction and merging bins. We ran *finaletoolkit delfi* with 1, 2, 4, 8, 16, 32, and 64 worker processes.

We used a fork of the original DELFI scripts (https://github.com/LudvigOlsen/delfi_scripts) due to the original DELFI implementation being highly *ad hoc* and difficult to use. This fork simplifies the command line interface, makes some optimizations on memory, and adds compatibility with BAM files aligned to GRCh38. This fork of DELFI scripts consists of 9 programs that are run in succession: *00-create_cytosine_file.r*, *00-filtered_regions.r*, *01-read_galp.r*, *02-fragment_gc.r*, *03.pre-create_bin_coordinates.r*, *03-bin_compartments.r*, *03.5-combine_bins.r*, *04-5mb_bins.r*, *06-gbm_full.r*. We ran this workflow using the “hg19” assembly option, chromosome lengths for GRCh37, and the provided gaps and filter files. We encountered a memory problem when running *01-read_galp.r* for the BH01 dataset. We had to modify the code using *reducebyYield* (from *GenomicFiles* v1.8.0 package in R), save the BAM file fragments in chunks of 1M fragments, and concatenate them, which took 95 Gb memory. It took ~110Gb memory to run *02-fragment_gc.*r and *03-bin_compartments.r* for the BH01 dataset. We were unable to run this code on GRCh38-aligned data to provide a fair time and memory benchmark. To compare the results, we ran *finaletoolkit delfi* again on GRCh37-aligned BH01.

Similar to WPS, there are small differences between the results generated by our tools and the results presented by Cristiano et al. This is due to differences in how fragments are filtered, with Cristiano et al using the R packages GenomicAlignments and GenomicRanges, while we filtered reads with our own code using pysam.

##### Benchmark end motifs and MDS

The source code was not released in the original publication (Jiang *et al.* 2020). We compared FinaleToolkit to Freefly (https://github.com/hellosunking/Freefly), an open source tool created by one of the co-authors of that publication. To benchmark Freefly, we acquired the FASTQ files for BH01 and ran the *freefly* command.

To benchmark FinaleToolkit, we ran *finaletoolkit interval-end motifs* on 10kb non-overlapped intervals or *finaletoolkit end motifs* genome-widely at BH01 dataset with a .2bit file for GRCh37 using 1, 2, 4, 8, 16, 32, and 64 worker processes. We then ran *finaletoolkit interval-mds* or *finaletoolkit mds* on the resulting tab-separated file to generate MDS data.

##### Benchmark Cleavage Profiles

The source code was not released in the original publication (Zhou *et al.* 2022). To benchmark FinaleToolkit, we ran *finaletoolkit cleavage-profile* on 100 kb intervals covering all autosomes with BH01 data using 1, 2, 4, 8, 16, 32, and 64  worker processes.

##### Benchmark Fragment Coverage

To benchmark FinaleToolkit, we ran *finaletoolkit coverage* on 100 kb intervals covering all autosomes with BH01 data using 1, 2, 4, 8, 16, 32, and 64 worker processes.

##### Benchmark Breakpoint motifs

The source code was not released in the original paper (Guo *et al.* 2022). To benchmark FinaleToolkit, we ran *finaletoolkit coverage* on 100 kb intervals covering all autosomes with BH01 data using 1, 2, 4, 8, 16, 32, and 64 worker processes.

#### Supplementary Figures

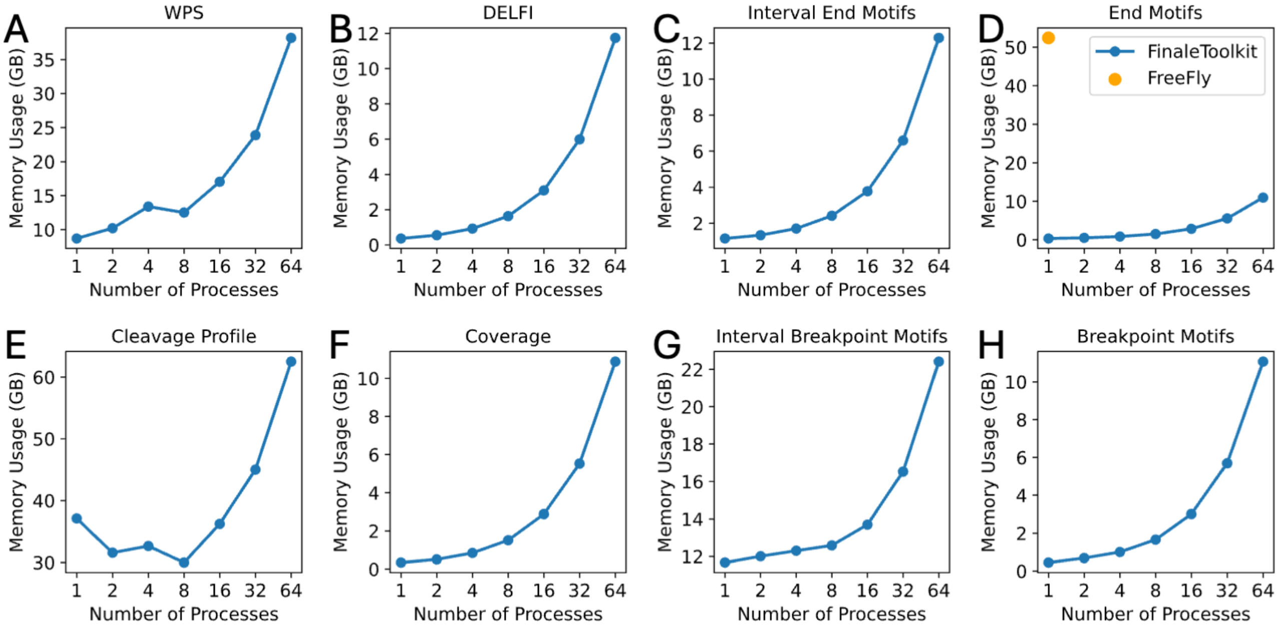


**Supplementary Figure 1. Memory usage of FinaleToolkit on BH01.** Memory cost to calculate (a) WPS, (b) cleavage ratio, (c) end motifs over intervals,  (d) genome-wide end motif, (e) DELFI, (f) fragment coverage, (g) breakpoint motifs over genome-wide intervals, and (h) breakpoint motifs without normalization with different numbers of processes.

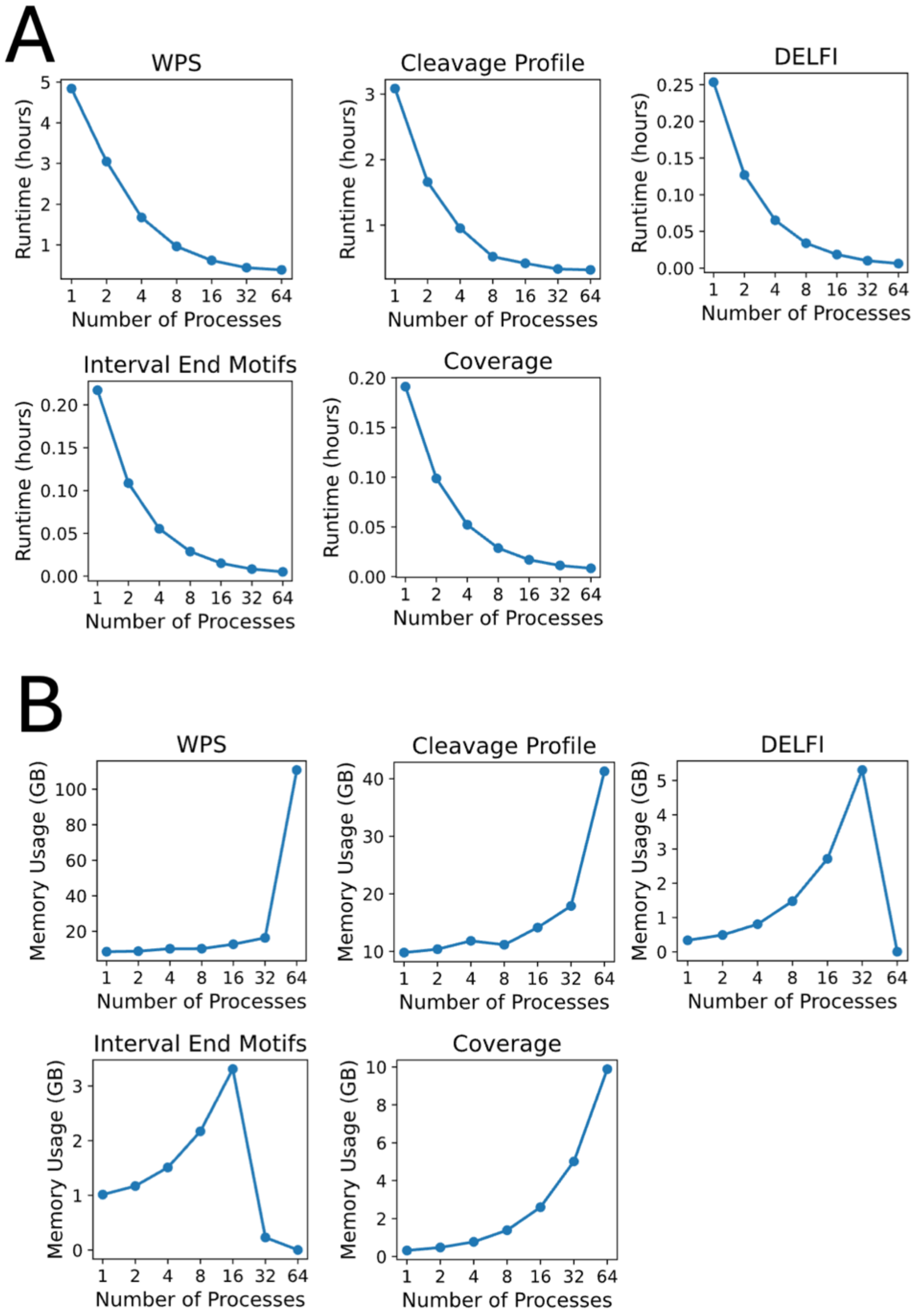


**Supplementary Figure 2. Performance of FinaleToolkit on low coverage cfDNA WGS data for select features.** We benchmarked the (a) wall time and (b) memory usage of FinaleToolkit using different numbers of CPU cores on H292 sample (Jiang *et al.* 2015) (~0.96X coverage).

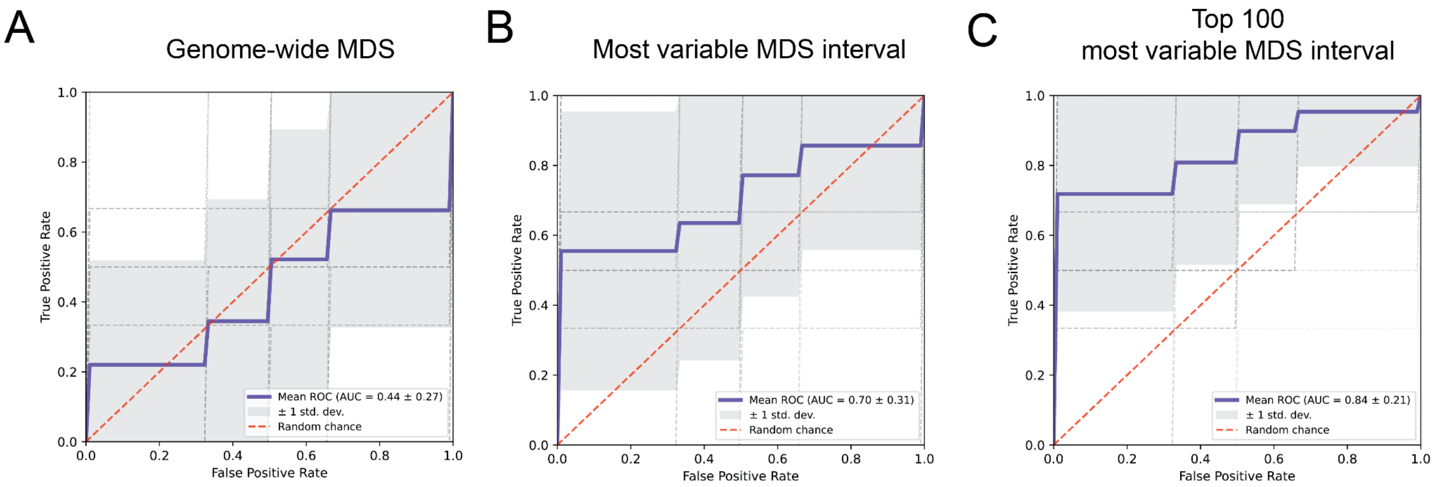


**Supplementary Figure 3. FinaleToolkit boosted the performance of cfDNA fragmentomics by genome-wide analysis of MDS over arbitrary genomic intervals.** The receiver operating characteristic (ROC) curve for distinguishing 25 early-stage breast cancer vs. 25 matched healthy controls by using (a) the genome-wide MDS score, (b) the most variable feature, and (c) the top 100 most variable features from MDS over 100kb non-overlapped intervals. The performance is calculated by applying logistic regression with l2 regularization at 10-fold cross-validation and repeated ten times. For a fair comparison, (a), (b), and (c) are compared with the same data split. The shade represents the standard deviation of ROC across repeated experiments.
